# Identifying different types of tumors in human brain, breast, and uterus according to their optical properties in the red to near-infrared region

**DOI:** 10.1101/2020.04.02.021568

**Authors:** Omnia Hamdy

## Abstract

Optical diagnosis techniques are gaining validity due to their competitive advantages including safety and functionality. These methods probe the tissue using light in the red to near-infrared region that shows relatively high penetration in biological tissues. The reflected light which is controlled by tissue absorption and scattering parameters can provide significant information about its pathology. In the present work, a method for identifying different types of tumors in human brain, breast and uterus based on their absorption and scattering characteristics in the red to near-infrared spectra is proposed. The classification method depends on studying the red to near-infrared light diffusion through the examined tissues using Monte-Carlo simulation and finite element solution of the light diffusion equation. The obtained tissue reflectance profiles, optical fluence rate distribution and absorbed faction in each tumor type show different characteristics along the selected wavelengths. The variation in these three features can be considered as an alternative approach of medical diagnosis or can assist with other conventional diagnosis methods.

## Introduction

Brain, breast and uterus are very vital and complex human body organs that have affected by many types of tumors with different characteristics and symptoms. Diagnosing brain tumors can be accomplished using different medical imaging techniques such as magnetic resonance imaging (MRI) and computer tomography (CT) scan or by a neurological exam [1]. While, X-ray mammography, MRI, breast ultrasonography and breast biopsies have been employed to diagnose breast tumors [2]. Although, MRI and CT can be used also to diagnose uterus tumors, transvaginal ultrasound and biopsy are the most common utilized techniques [3]. However, many of these techniques have their limitations. Besides, the use of ionizing radiation in the diagnosing process can even develop tumors in the healthy tissues [4]. On the other hand, optical techniques are considered very safe and functional diagnosing tools.

When light in red to near infrared region interacts with biological tissues it diffusely propagates due to the inhomogeneity and multiple scattering characteristics of tissues [5,6]. A part of the penetrated light might be absorbed by specific tissue chromophores, or it can change its direction inside the tissue layers by scattering [7,8]. Otherwise, some of this light leaves the tissue either by transmission or reflection. The optical absorption and scattering properties of any biological tissue can be utilized as indictors for tissue health [9] as they are directly related to tissue chromophores concentrations including water contents, oxygenation of hemoglobin and melanin concentration [10,11]. Moreover, tissue diffuse reflectance is considered as a finger print for each tissue type and condition as it changes upon appearing any abnormality in the tissue [12]. Therefore, both tissue optical parameters and diffuse reflectance profile can powerfully assist in the process of medical diagnosis and therapy monitoring [6,13–15]. Visible and near infrared diffuse optical spectroscopic techniques have been widely used in breast cancer diagnosis process [16,17] and in differentiating malignant tumor tissue from benign tumor and normal breast tissues [18,19].

Tissue diffuse reflectance and transmittance are the main measurements required to determine the optical parameters of any type of biological tissue, they can be experimentally measured by either integrating spheres [20] or distant photo-detectors [21]. Based on the reflection and transmission collected data, tissue absorption coefficient µa, scattering coefficient µs and anisotropy factor g can be estimated using various analytical approaches referred as indirect or inverse methods such as Kubelka-Munk mathematical model [22], inverse Monte-Carlo [23] and inverse adding-doubling iterative method [24]. Other techniques referred as forward or direct methods that use the known optical parameters values to obtain the diffuse reflectance and transmission profiles of the studied tissue based on the radiative transport theory of light propagation [25]. However, Monte-Carlo (MC) simulation is an alternative numerical method that is widely used to simulate the light propagation in multiple scattering medium such as biological tissues [26]. It was utilized to study the light transmission properties in breast tissue in order to improve the imaging systems considered to detect breast cancer [27,28] and to detect different breast diseases based on the light propagation in the breast tissue [29,30]. Moreover, MC method was implemented to estimate the x-ray absorption in the brain tumor during different radiotherapy clinical procedures [31] and in stereotactic radiotherapy [32]. It has been also employed to improve the detection process of brain tumors in MRI images [33]. In addition, MC method played a useful role in enhancing the detection of cervical dysplasia and cancer [34,35].

In this work, different types of tumors in human brain, breast and uterus have been identified according to its optical absorption and scattering properties in the optical window (600-900nm). The classification method depended on three features; the spatially resolved diffuse reflectance, the optical fluence rate and the tissue absorbed fraction. The fluence rate distribution at the tissue surface was obtained using finite element solution of the diffusion equation, while, the diffuse reflectance profiles and absorbed fraction were acquired using Monte-Carlo simulation for light propagation in biological tissues.

## Methods

### Light Transport in Biological Tissue

Recent advances in describing laser transfer energy in biological tissues are based on transport theory of light propagation that utilizes the radiative transport equation RTE to represent this propagation [36]. The radiative transport equation can be written as. [7]

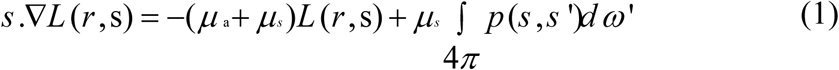

where L(r,s) is the radiance of light at position r traveling in a direction of the unit vector s and 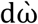 is the differential solid angle in the direction 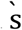. µa is the tissue absorption coefficient and µs is the scattering coefficient.

When a laser beam is incident on a turbid medium as biological tissue, the radiance can be divided into a coherent and diffuse term. The diffuse radiance is the main problem which transport theory has to deal with, since scattered photons do not follow a determined path. Therefore, many approximations and statistical approaches were introduced to the transport equation depending on the value of the albedo of the medium [37].

Using radiative equation to calculate the light distribution in tissue requires the knowledge of the absorption and scattering coefficients as well as the phase function. In order to estimate these parameters, one must have a solution of the radiative transport equation. The diffusion equation (2) is an approximation to the radative transport equation in which the radiance is assumed to be nearly isotropic, it is a partial differential equation that is significantly easier to be solved than the radiative transport equation [38].

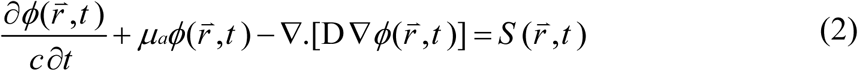

where the constant 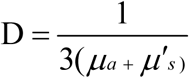 is the diffusion coefficient and *µ*′_*s*_ = (1−*g*) *µ* _*s*_ is the reduced scattering coefficient and g is the anisotropy factor. 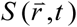 represents the source term and 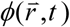 is the fluence rate.

In this study, the diffusion equation was solved using the finite element method [39,40] under the environment of COMSOL Multiphysics 5.2 software [41]. Helmholtz Equation (4) in COMSOL is employed to model the diffusion equation presented in (3) in the steady state.

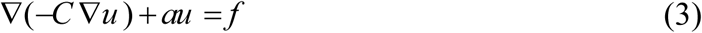

Comparing the parameters in equation (3) with those in (2) we get, *u = ϕ, C* = *D, a* = *μ*_*a*_, and *f* = *S*

A 1 cm circular model was created with a point source at the center of the model as presented in Fig. 1(a). The implemented mesh (Figure 1(b)) has element size ranges from 0.000057 to 0.02 cm.

**Fig. 1.**
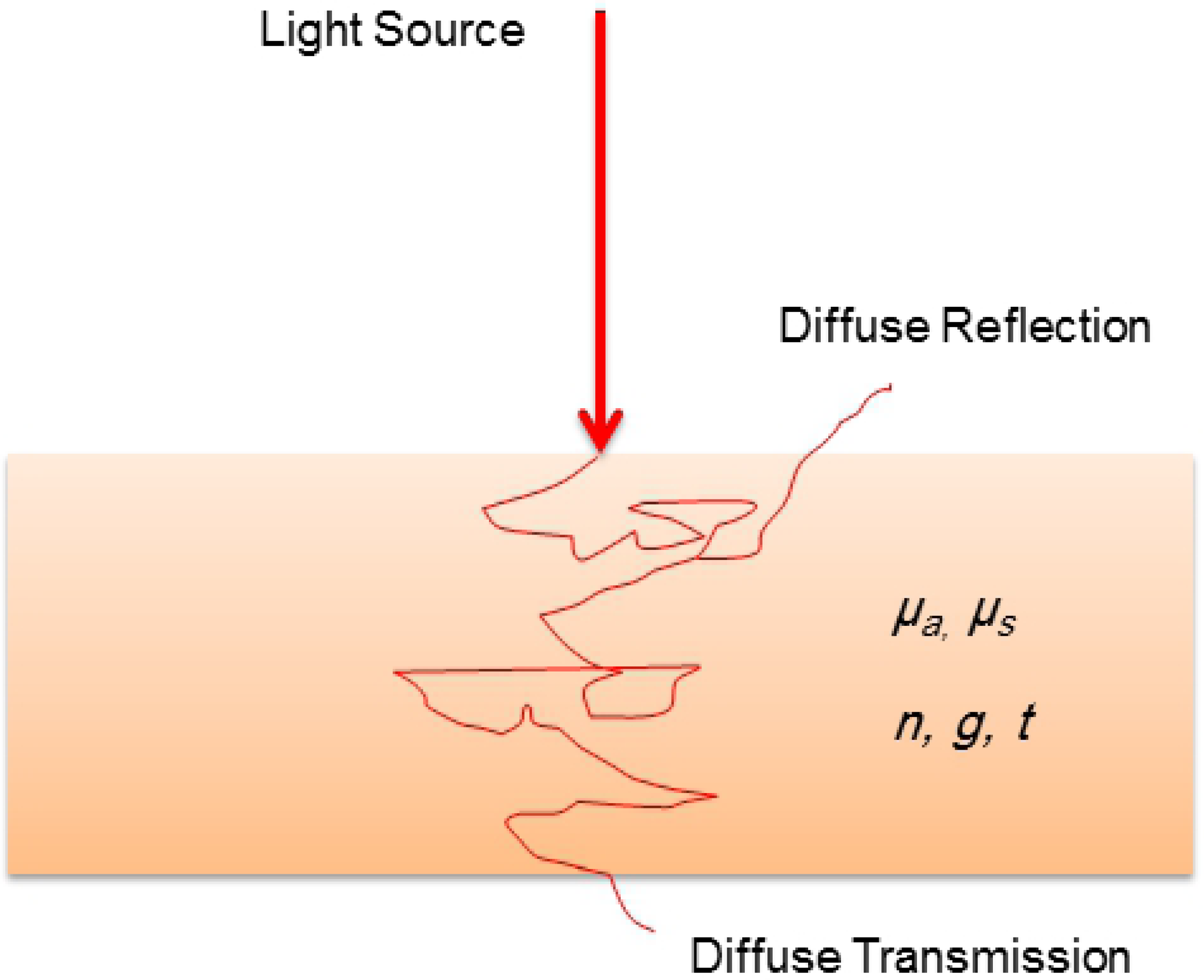
The implemented finite element (a) model, (b) mesh

Based on the predefined finite element model, the fluence rate distribution at the surface of the various selected tissues was obtained.

### Monte-Carlo simulation for light propagation

Monte-Carlo simulation for light propagation in biological tissues (MCML) is a numerical approach that is widely used to simulate the light propagation in turbid medium [42,43]. The simulation is designed to deal with either single layer or multi-layers tissue models. However, each tissue layer has to be defined with its refractive index n, absorption coefficient µ_a_, scattering coefficient µ_s_, anisotropy g and thickness t [44] as illustrated in Fig. 2.

**Fig. 2.**
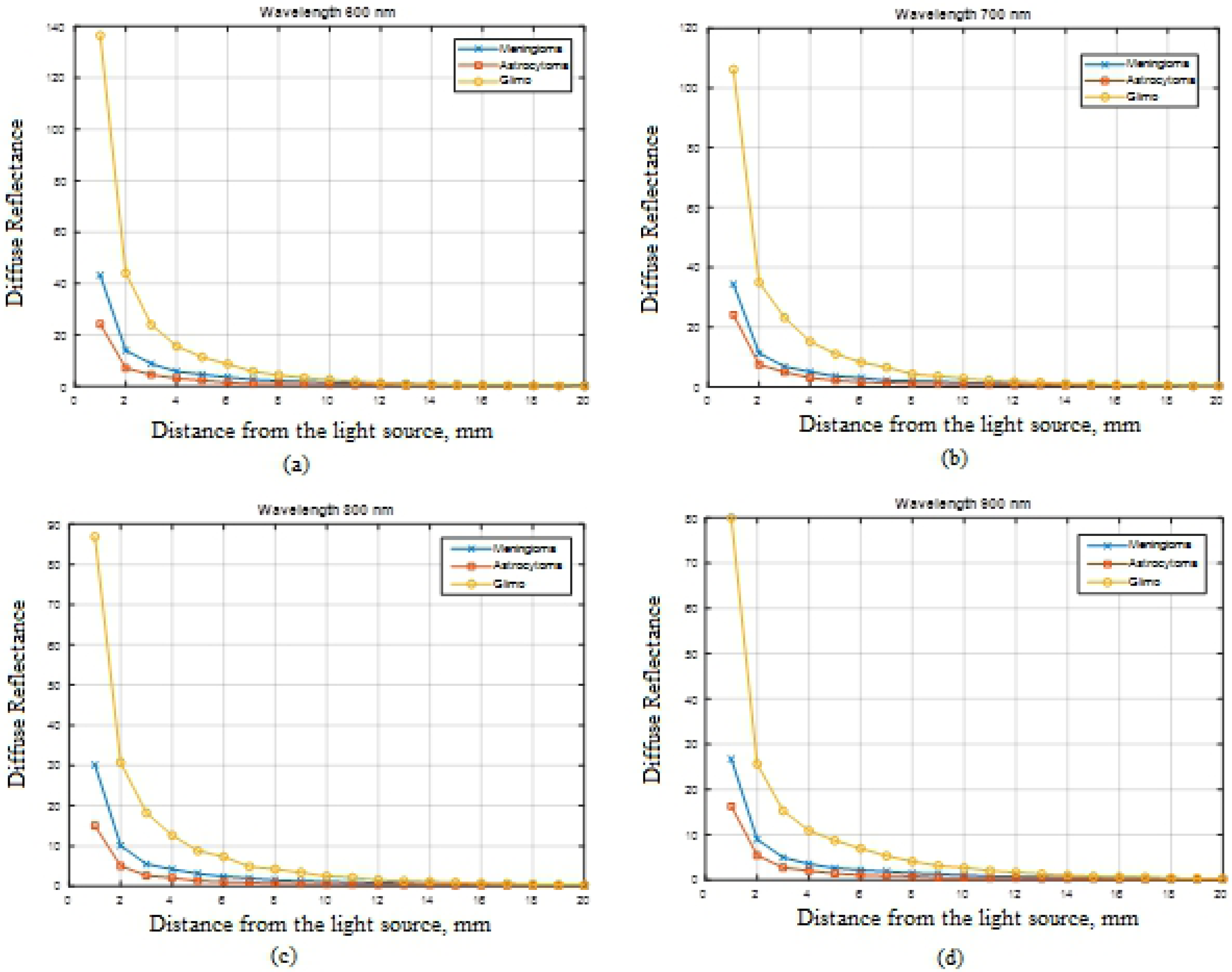
Monte-Carlo Simulation model

### Selected tissue samples

In this study, common types of tumors (benign and malignant) in human brain, breast and uterus with previously published optical absorption and scattering parameters [25] have been examined. The selected tissue types are; meningioma, astrocytoma and glioma brain tumors, fibrocystic, fibroadenoma and ductal carcinoma of breast and finally, myometrium and leiomyoma uterus tumors. Table 1 presents the optical parameters utilized in this implantation.

**Table 1.** The utilized tissues optical parameters [25].

| Tissue | Wavelength (nm) | $\mu_a$ (cm<sup>-1</sup>) | $\mu_s$ (cm<sup>-1</sup>) | $g$ |
| --- | --- | --- | --- | --- |
| Meningioma | 600 | 0.67 | 175.0 | 0.953 |
|  | 700 | 0.30 | 154.9 | 0.956 |
|  | 800 | 0.23 | 138.9 | 0.959 |
|  | 900 | 0.22 | 125.9 | 0.958 |
| Astrocytoma | 600 | 1.19 | 130.4 | 0.962 |
|  | 700 | 0.41 | 112.0 | 0.960 |
|  | 800 | 0.50 | 96.94 | 0.967 |
|  | 900 | 0.32 | 86.62 | 0.963 |
| **Glioma** | 600 | 2.35 | 227.4 | 0.9 |
|  | 700 | 1.42 | 183.4 | 0.9 |
|  | 800 | 1.35 | 155.8 | 0.9 |
|  | 900 | 1.40 | 144.7 | 0.9 |
| **Fibrocystic** | 600 | 0.47 | 623.5 | 0.977 |
|  | 700 | 0.24 | 568.0 | 0.978 |
|  | 800 | 0.28 | 548.8 | 0.981 |
|  | 900 | 0.39 | 536.5 | 0.981 |
| **Fibroadenoma** | 600 | 1.76 | 492.6 | 0.979 |
|  | 700 | 0.53 | 438.4 | 0.982 |
|  | 800 | 0.34 | 384.0 | 0.983 |
|  | 900 | 0.79 | 327.3 | 0.983 |
| **Ductal Carcinoma** | 600 | 1.61 | 337.5 | 0.958 |
|  | 700 | 0.44 | 277.4 | 0.961 |
|  | 800 | 0.34 | 233.0 | 0.962 |
|  | 900 | 0.45 | 181.2 | 0.957 |
| **Myometrium** | 610 | 0.45 | 133.0 | 0.9 |
|  | 700 | 0.19 | 114.7 | 0.9 |
|  | 800 | 0.10 | 90.8 | 0.9 |
|  | 900 | 0.10 | 73.2 | 0.9 |
| **Leiomyoma** | 600 | 0.15 | 109.9 | 0.9 |
|  | 700 | 0.07 | 87.3 | 0.9 |
|  | 800 | 0.03 | 67.5 | 0.9 |
|  | 900 | 0.03 | 54.7 | 0.9 |

## Results and discussion

### Diffuse reflectance profiles

Based on Monte-Carlo simulation for light propagation in biological tissues, the spatially resolved steady state diffuse reflectance profiles for different tumor types were obtained. Fig. 3 illustrates the variation in the reflectance profiles between meningioma, astrocytoma and glioma.

**Fig. 3.**
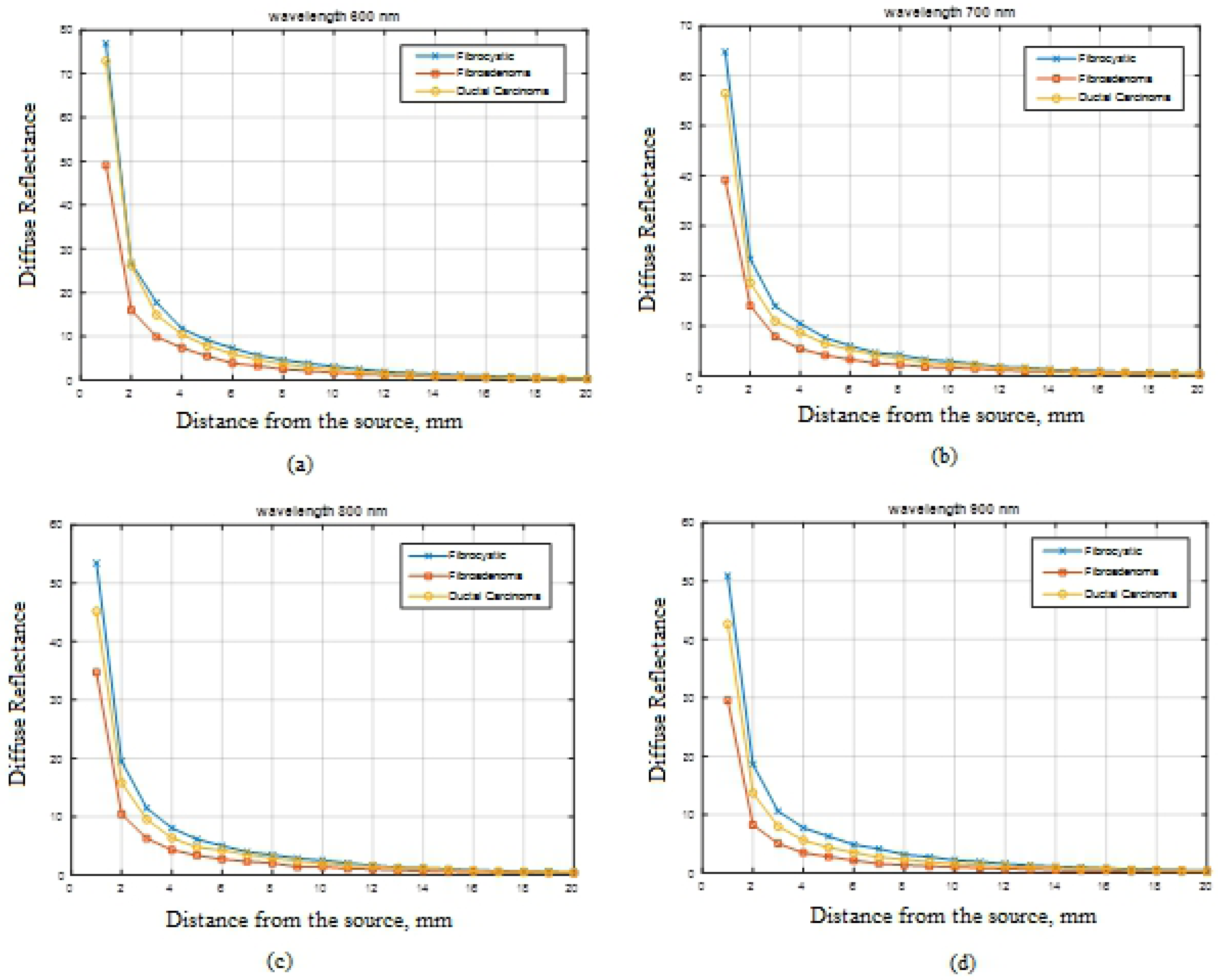
Spatial distribution of light in brain tumors at different wavelengths

Meningioma showed maximum reflectance values over the all implemented wavelengths followed by astrocytoma then glioma. The reflectance behavior is almost the same at every wavelength. While in case of breast tumors, fibrocystic and fibroadenoma are showing very close values at 600 and 700 nm, however, the reflection profile begins to deviate at 800 and 900 nm as presented in Fig. 4.

**Fig. 4.**
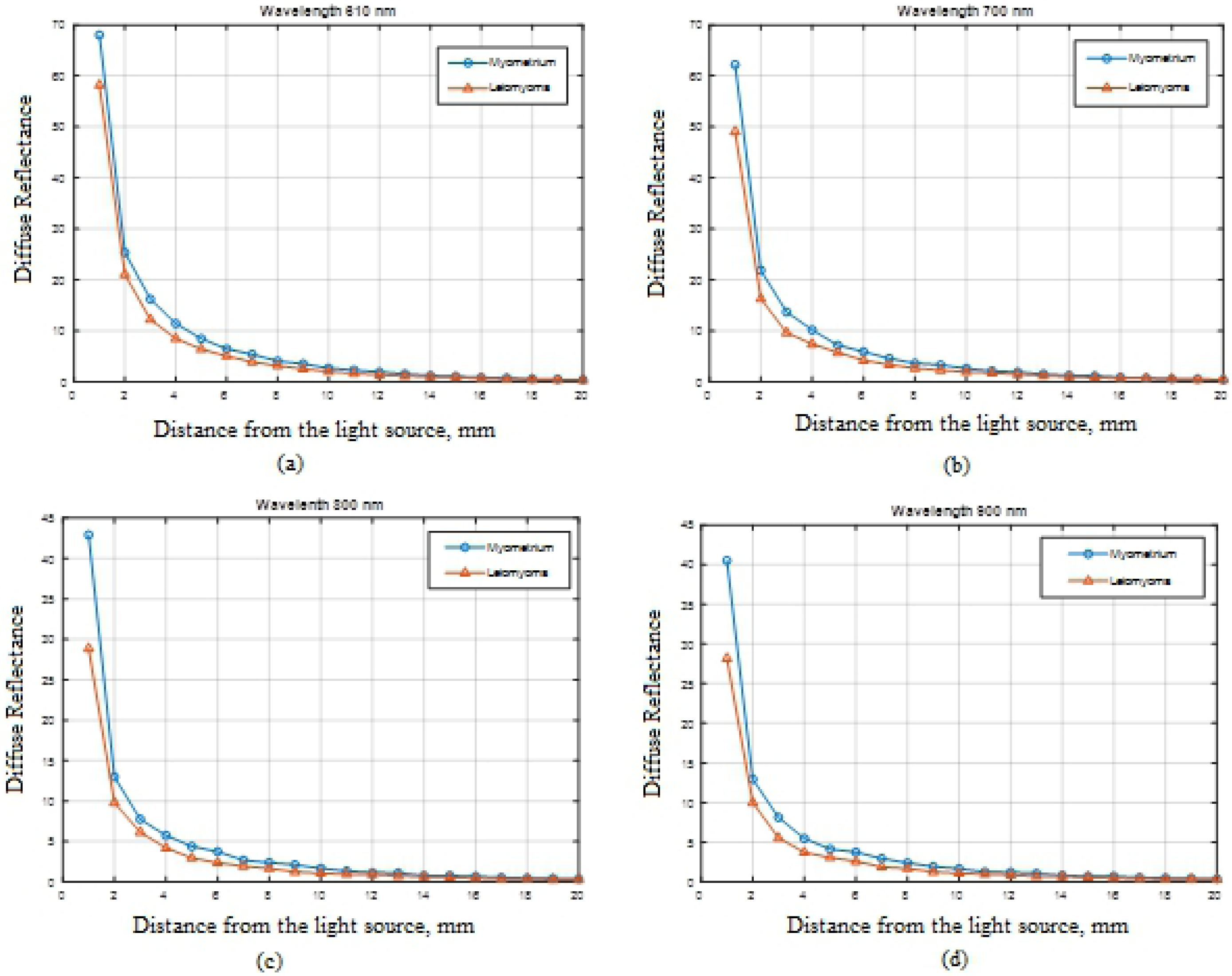
Spatial distribution of light in breast tumors at different wavelengths

Ductal carcinoma can be clearly differentiated from other types of the examined tumors as it always has the minimum reflectance values at every wavelength. Regarding uterus tumors, myometrium shows higher values than leiomyoma over the four selected wavelengths as illustrated in Fig. 5.

**Fig. 5.**
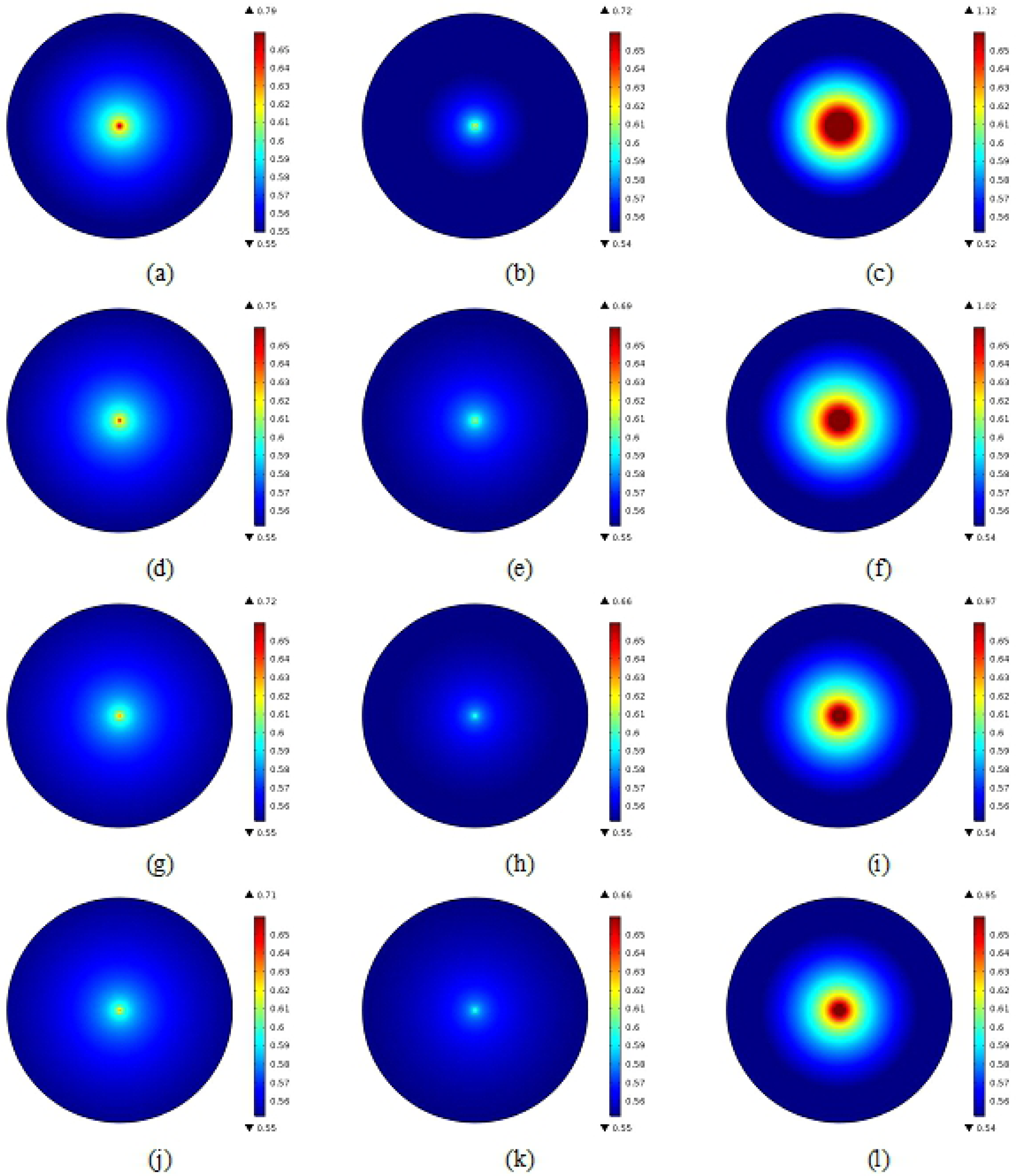
Spatial distribution of light in uterus tumors at different wavelengths

### Fluence rate distribution

The optical fluence rate distribution at the surface of the examined tumors has been visualized using finite element method. Fig. 6 illustrates the distribution in meningioma, astrocytoma ad glioma at the selected wavelengths. Glioma recorded higher fluence rate distribution at each wavelength followed by meningioma then astrocytoma. Therefore, the obtained images can be significantly used to distinguish between these types of tumors.

**Fig. 6.**
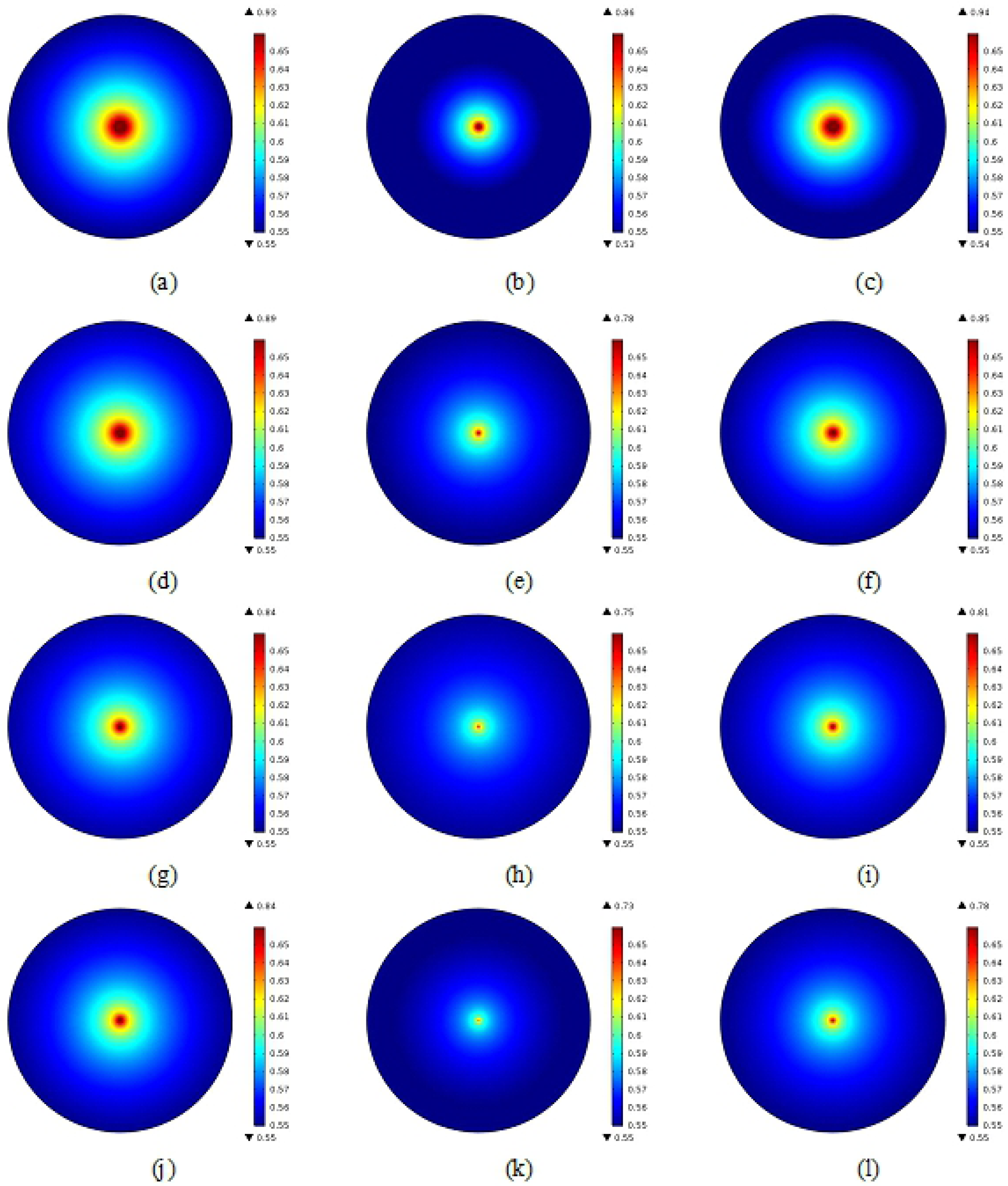
(a) Meningioma at 600 nm, (b) Astrocytoma at 600 nm, (c) Glioma at 600 nm, (d) Meningioma at 700 nm, © Astrocytoma at 700 nm, (f) Glioma at 700 nm, (g) Meningioma at 800 nm, (h) Astrocytoma a at 800 nm, (i) Glioma at 800 nm, (j) Meningioma at 900 nm, (k) Astrocytoma at 900 nm, (l) Glioma at 900 nm,

Analyzing the fluence distribution in the breast tumors revealed that, fibroadenoma gives the minimum distribution at all implemented wavelengths as presented in Fig. 7. Fibrocystic and ductal carcinoma show almost the same distribution at 600 nm, while from 700 to 900 nm, ductal carcinoma has lower values compared with fibrocystic type.

**Fig. 7.**
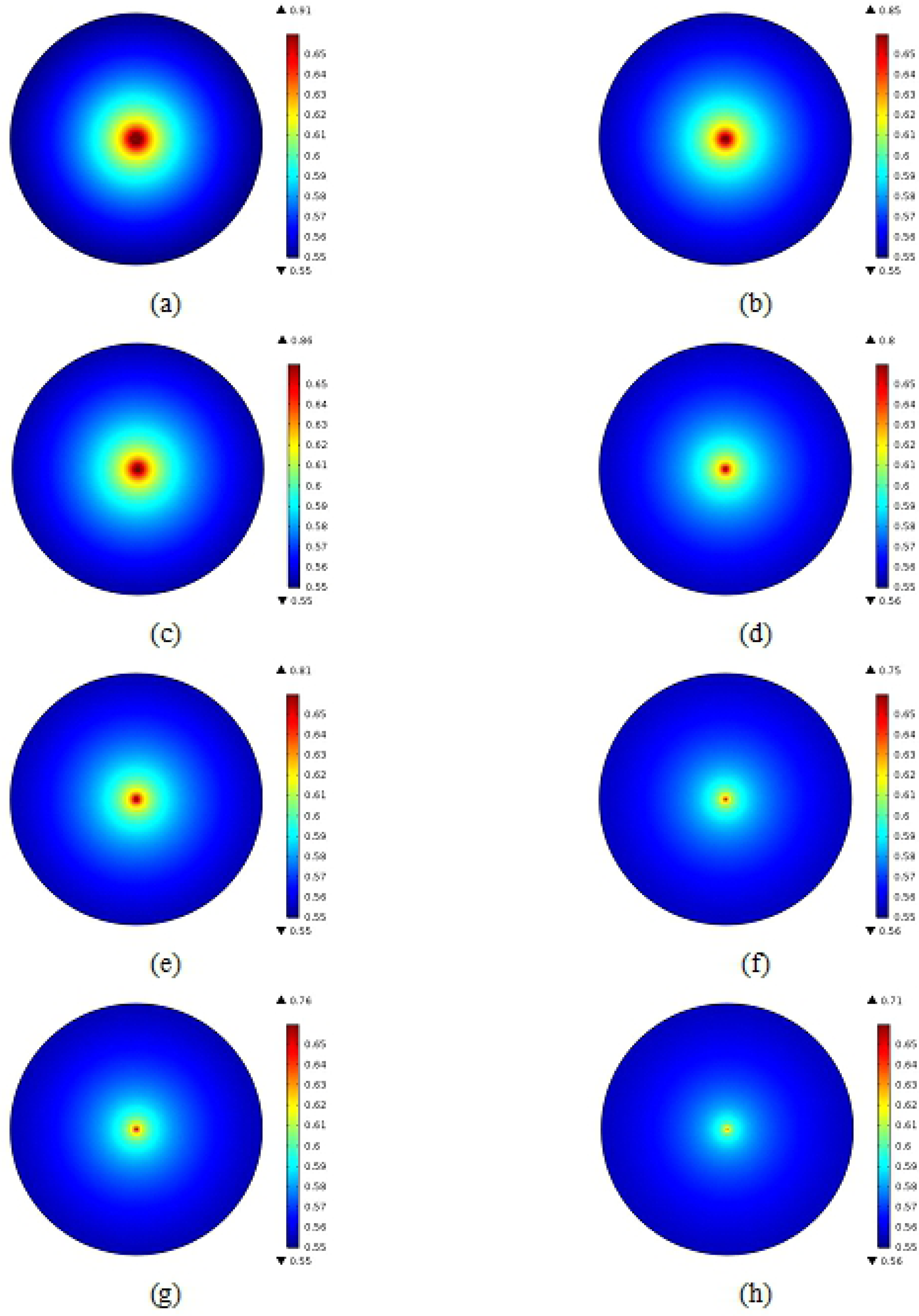
(a) Fibrocystic at 600 nm, (b) Fibroadenoma at 600 nm, (c) Ductal Carcinoma at 600 nm, (d) Fibrocystic at 700 nm, © Fibroadenoma at 700 nm, (f) Ductal Carcinoma at 700 nm, (g) Fibrocystic at 800 nm, (h) Fibroadenoma at 800 nm, (i) Ductal Carcinoma at 800 nm, (j) Fibrocystic at 900 nm, (k) Fibroadenoma at 900 nm, (l) Ductal Carcinoma at 900 nm,

In case of uterus tumors, myometrium shows higher fluence rate than Leiomyoma at all studied wavelengths as presented in Fig. 8. The obtained images are also useful in discriminating these two types of uterus tumors.

**Figure 8.**
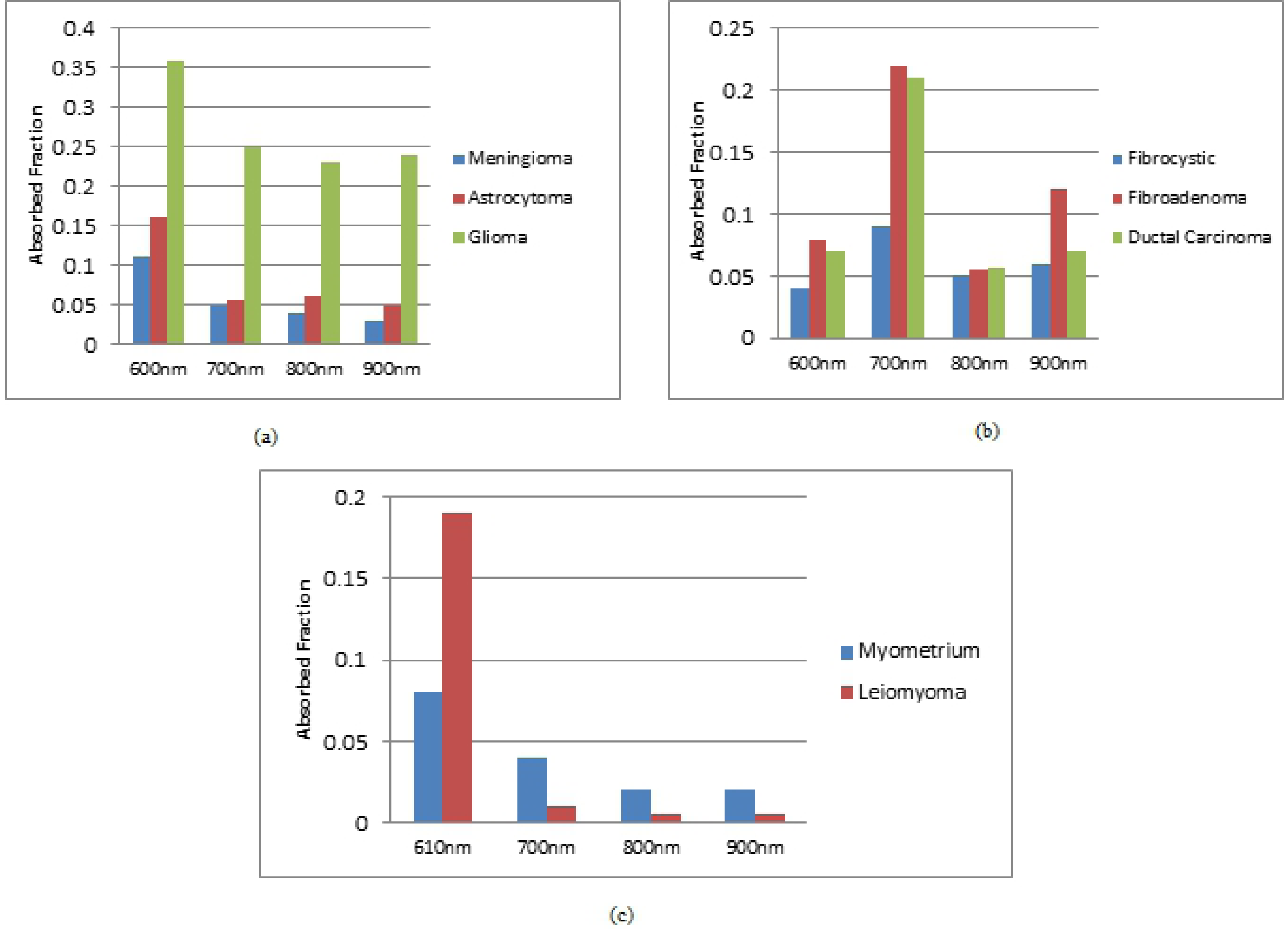
The distribution of fluence rate of (a) Myometrium at 610 nm, (b) Leiomyoma at 610 nm (c) Myometrium at 700 nm, (d) Leiomyoma at 700 nm © Myometrium at 800 nm, (f) Leiomyoma at 800 nm (g) Myometrium at 900 nm, (h) Leiomyoma at 900 nm

### Tissue absorbed fraction

MCML simulation output can provide information about he absorbed fraction in the studied sample. Fig. 9 (a), (b) and (c) Illustrate the variation in the absorbed fraction in brain, breast and uterus tumors respectively.

**Fig. 9.**
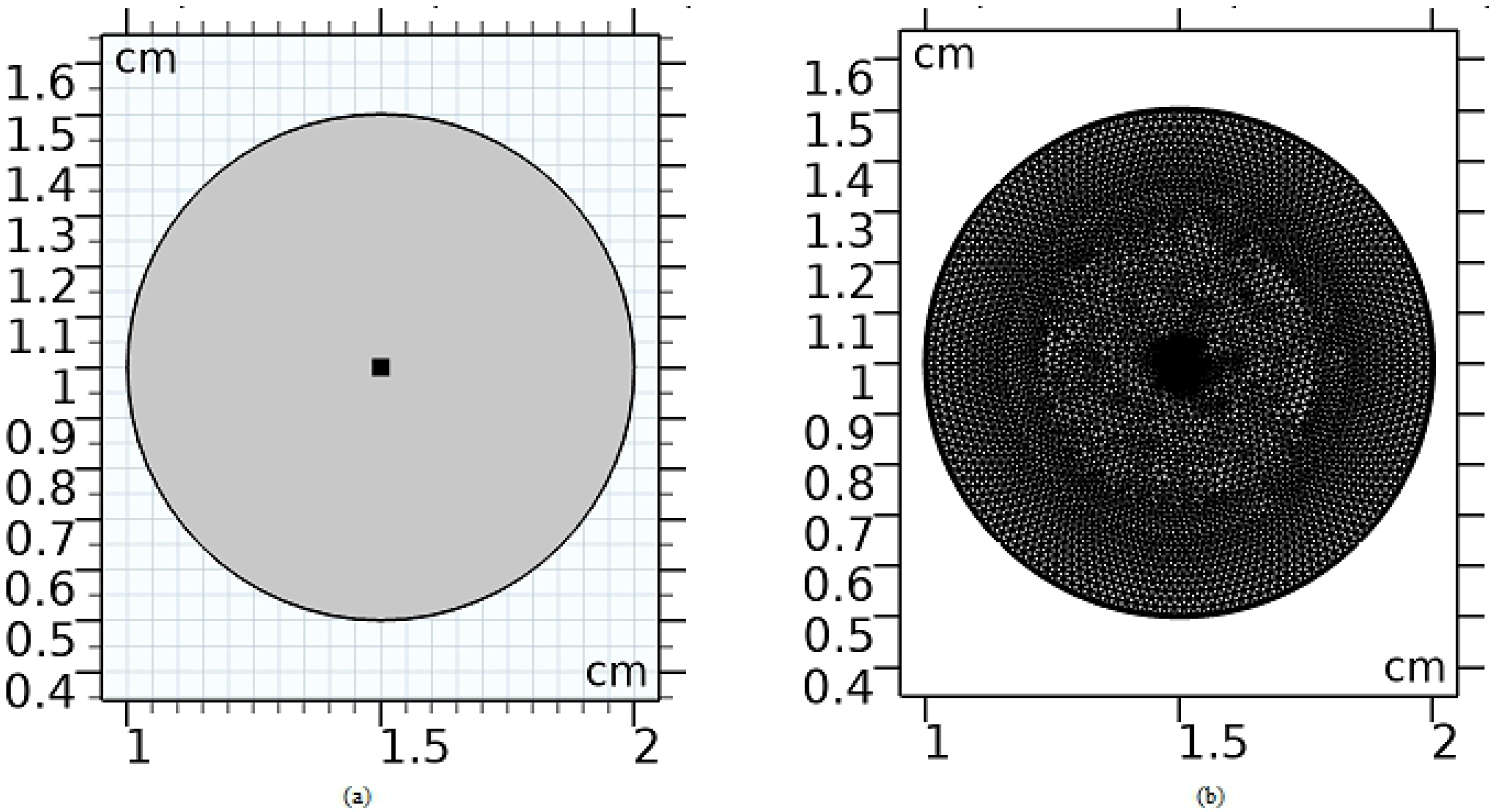
The variation in tissue absorbed fraction of (a) brain tumors, (b) in breast tumors, (c) uterus tumors

Meningioma shows higher absorbed fraction than astrocytoma and glioma at all wavelengths. While in breast tumors, fibrocystic records lower absorbed fraction than fibroadenoma and ductal carcinoma. Myometrium shows the highest absorbed fraction at 700, 800, and 900 nm, while at 610 nm, leiomyoma recorded much higher value than myometrium.

The diffuse reflectance curves presented in Figs. 3, 4 and 5 provide a different profile at each studied type of tumor. These profiles are changing as a result of the variation in absorption and scattering parameters between each tumor at every wavelength. These parameters have also a significant effect on the tissue absorbed fraction and fluence rate distribution at the tumor tissue surface. Although meningioma has lower absorption and scattering coefficients, it shows higher absorbed fraction and reflectance values but less fluence rate than glioma. Fibrocystic tissue has lower absorption but higher scattering coefficients, thus, it has the minimum absorbed fraction and maximum fluence rate. Myometrium shows higher reflectance and fluence rate values than leiomyoma as it has higher absorption and scattering coefficients as shown in table 1, therefore myometrium recorded also higher absorbed fraction.

## Conclusion

In conclusion, a medical diagnosis approach based on tissue diffuse reflection and fluence rate distribution is presented and investigated. The presented investigations depended on Monte-Carlo simulation and the finite element solution of the diffusion equation. The proposed results have demonstrated the ability of diffuse reflectance profiles, fluence rate distribution and tissue absorbed fraction monitoring to identify different types of tumors in human brain, breast and uterus. The difference in the reflection profiles, absorbed fraction values and fluence rate images between each studied tissue provides a significant criterion for identifying various types of tumors.

